# DERM-Array: An Open-Source Platform for Spatially Resolved Tumor Immunology

**DOI:** 10.64898/2026.11.25.769288

**Authors:** Melanie Meyer, Shaina Hall, Kelsey Furiya, Runab Dixit, Ashwika Mattam, Jason McBane, Raymond Ruff, Sawyer Sarcot, Jason P. Price, Catherine B. Carbone

## Abstract

Tumor arrays allow researchers to study dozens of tumors within a single mouse to investigate how local microenvironments influence tumor growth. Here we describe dermal engraftment of regional microtumors-array (DERM-Array), an open-source method that uses laser implantation to generate tumor arrays in the skin of a mouse ear. With DERM-Array, we demonstrate longitudinal *in vivo* imaging across multiple tumor models to characterize tumor growth, spatial immune organization, and immunotherapy response.

## Main Text

Tumor arrays are a tool to investigate how local tumor microenvironments influence tumor growth and treatment response by comparing dozens of spatially distinct tumors exposed to the same systemic environment. These models enable both real-time imaging of microenvironment dynamics and longitudinal tracking of individual lesions over several weeks of tumor growth and treatment. The previously described skin tumor array by micro-poration (STAMP) platform demonstrated the utility of this approach, revealing that genetically identical tumors can develop distinct immune microenvironments and undergo different fates within the same mouse (Ortiz-Muñoz et al., 2023). The original study demonstrated that early T cell infiltration predicts tumor rejection and identified a fibroblast-myeloid cell signaling axis that promotes T cell recruitment. Subsequent work showed that intratumoral cDC1-T cell interactions provide local costimulation that differentiates rejected tumors from those that persist despite T cell infiltration (Carbone et al., 2026). Together, these studies establish tumor arrays as a powerful experimental platform for identifying the cellular interactions within tumor microenvironments that promote or prevent tumor rejection.

Although tumor arrays have demonstrated considerable scientific value, their broader adoption has been limited by two practical barriers. First, previous studies demonstrated the biological utility of tumor arrays but did not provide sufficient technical guidance for other laboratories to reproduce the method independently. Second, the original implementation relied on an expensive laser system whose manufacturer has discontinued production of the device and its consumables (Ortiz-Muñoz et al., 2023; Carbone et al., 2026). To address these barriers, we developed dermal engraftment of regional microtumors-array (DERM-Array), an open-source method that introduces a new approach to laser implantation using inexpensive, commercially available programmable diode laser engravers (**Fig. 1A**). Here, we provide detailed experimental protocols and validation benchmarks for reproducibly generating tumor arrays and performing longitudinal imaging and downstream analyses.

**Figure 1.**
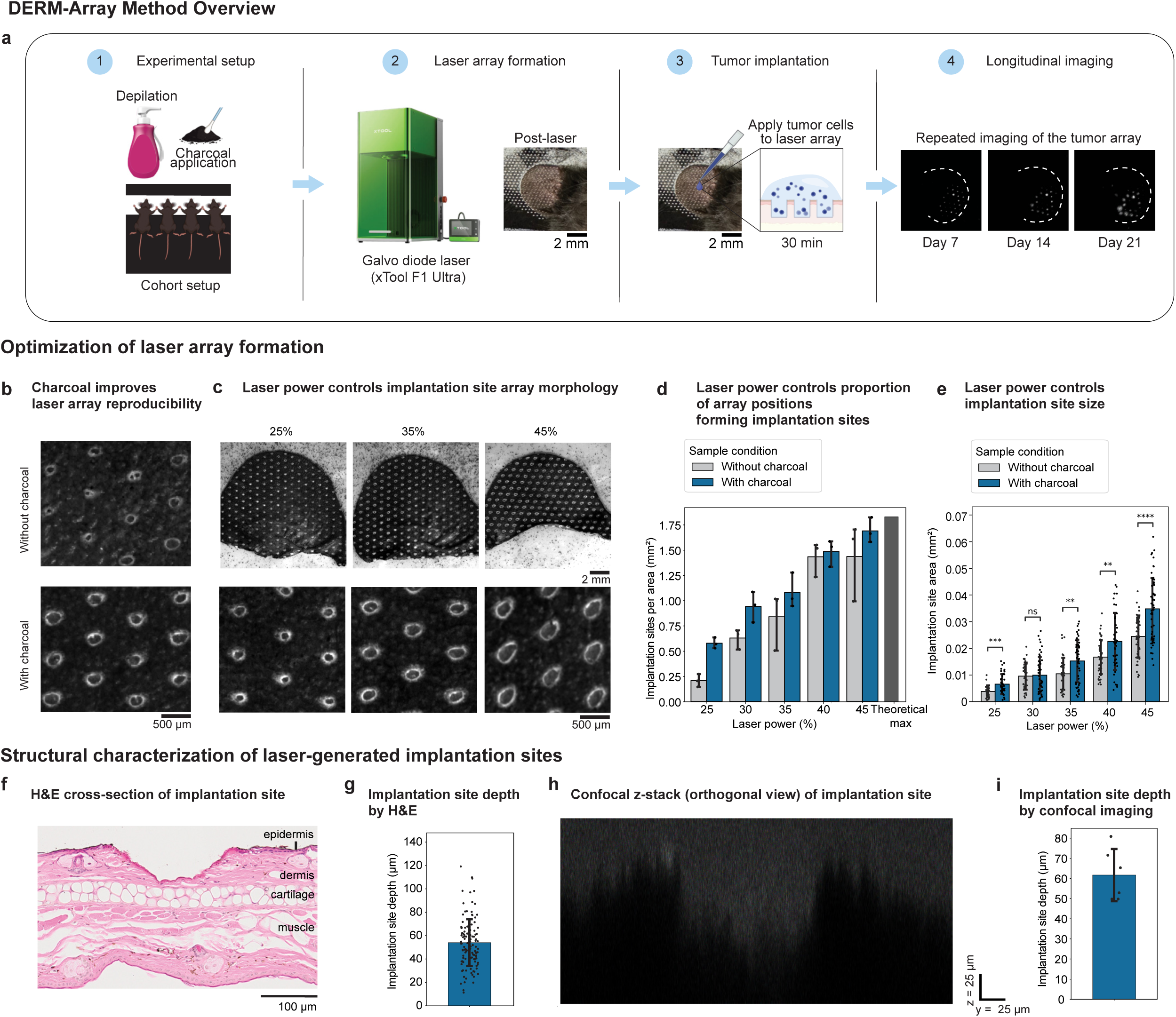
DERM-Array adapts chromophore-assisted laser ablation to generate arrayed tumor implantation sites. **a.** Schematic of the DERM-Array procedure showing ear depilation and charcoal application, laser array formation using the xTool F1 Ultra, tumor cell implantation and longitudinal fluorescence imaging. Scale bar, 2 mm. **b.** Representative brightfield images of implantation sites generated with or without charcoal application. Scale bar, 500 µm. **c.** Representative brightfield images of PFA-fixed ear pinnae with implantation sites generated at 25%, 35% and 45% laser power (5, 7 and 9 Watts). Scale bars, 2 mm (top) and 500 µm (bottom). **d,e.** Quantification of implantation site density (**d**) and area (**e**) in **c** (n = 3 ears per condition; 16–71 sites per condition). **f.** Representative H&E-stained cross-section of a laser-generated implantation site. Scale bar, 100 µm. **g**. Quantification of implantation site depth from H&E-stained sections in f (n = 125 sites from 5 ears) **h.** Representative orthogonal reconstruction of a confocal z-stack showing implantation site morphology on fresh tissue. Scale bar, 25 µm. **i.** Quantification of implantation site depth from confocal z-stacks in **h** (n = 5 sites from 3 ears). Each point in **d**, **e**, **g** and **i** represents an individual implantation site. Bars represent mean and error bars indicate standard deviation. Statistical analysis was performed using a two-tailed Mann–Whitney U-test.

Tumor arrays are generated by using a laser to create a grid of small implantation sites in the skin of a mouse ear, followed by topical application of tumor cells that engraft within these sites (**Fig. 1A**). The original STAMP method used an Er:YAG laser, which is strongly absorbed by water in the skin and produces localized tissue ablation (Ortiz-Muñoz et al., 2023; Carbone et al., 2026). To enable the use of more widely available lasers, we developed an alternative approach in which an exogenous chromophore is applied to the skin surface to absorb laser light and generate reproducible implantation sites. We implemented this method using a commercially available, programmable galvo-driven diode laser engraver (xTool F1 Ultra).

Unlike the manually positioned Er:YAG system, the new device uses software-controlled scanning to define implantation site locations, power, and dwell time, enabling simultaneous preparation of arrays across 3-5 mice (**Supplementary Fig. 1A,B**). Built-in safety features and a simple calibration procedure further support reproducible implementation (**Supplementary Fig. 1C and Supplementary Protocol).**

We next optimized skin preparation and laser settings to generate reproducible implantation sites. Treating ears with a depilatory cream and applying a thin layer of charcoal powder to the ear surface improved the uniformity of implantation sites by providing consistent absorption of visible laser light (**Fig. 1A,B**). Systematic laser power titrations established conditions that reliably generated implantation sites with regular dimensions (**Fig. 1C-E**). Histological and confocal characterization confirmed the formation of discrete implantation sites with depths of approximately 50-60 µm (**Fig. 1F-I**). Comparison with STAMP revealed similar implantation site depths but modest differences in diameter (**Supplementary Fig. 2**). Across a range of laser powers (30-40%), microtumor engraftment, persistence, and growth remained comparable, providing a practical operating range for array generation (**Supplementary Fig. 3**). To characterize the tissue response to laser injury, we performed bulk RNA sequencing of treated skin with and without tumor cells. Injury-associated inflammatory signatures, including neutrophil/IL-1 and interferon-response programs, increased during the first three days and declined toward baseline by day 14 (**Supplementary Fig. 4**).

Following laser ablation, tumor cells are applied to the ear surface and allowed to engraft for approximately 30 minutes (**Supplementary Protocol**). Individual microtumors become visible within seven days and can be imaged longitudinally for at least three weeks using a conventional fluorescence stereomicroscope (**Fig. 2A**). We established quantitative workflows for measuring implantation efficiency (**Fig. 2B**), microtumor persistence and rejection (**Fig. 2C,D**), and individual microtumor growth trajectories (**Fig. 2E-G; Supplementary Note 2**). These analyses capture heterogeneous tumor behaviors, including progression, stable growth, regression, and complete rejection. Direct comparison of B16F10-EGFP arrays generated using DERM-Array and STAMP revealed no statistically significant differences in microtumor engraftment, persistence, or growth, supporting the biological performance of the new implantation method (**Supplementary Fig. 5**).

**Figure 2.**
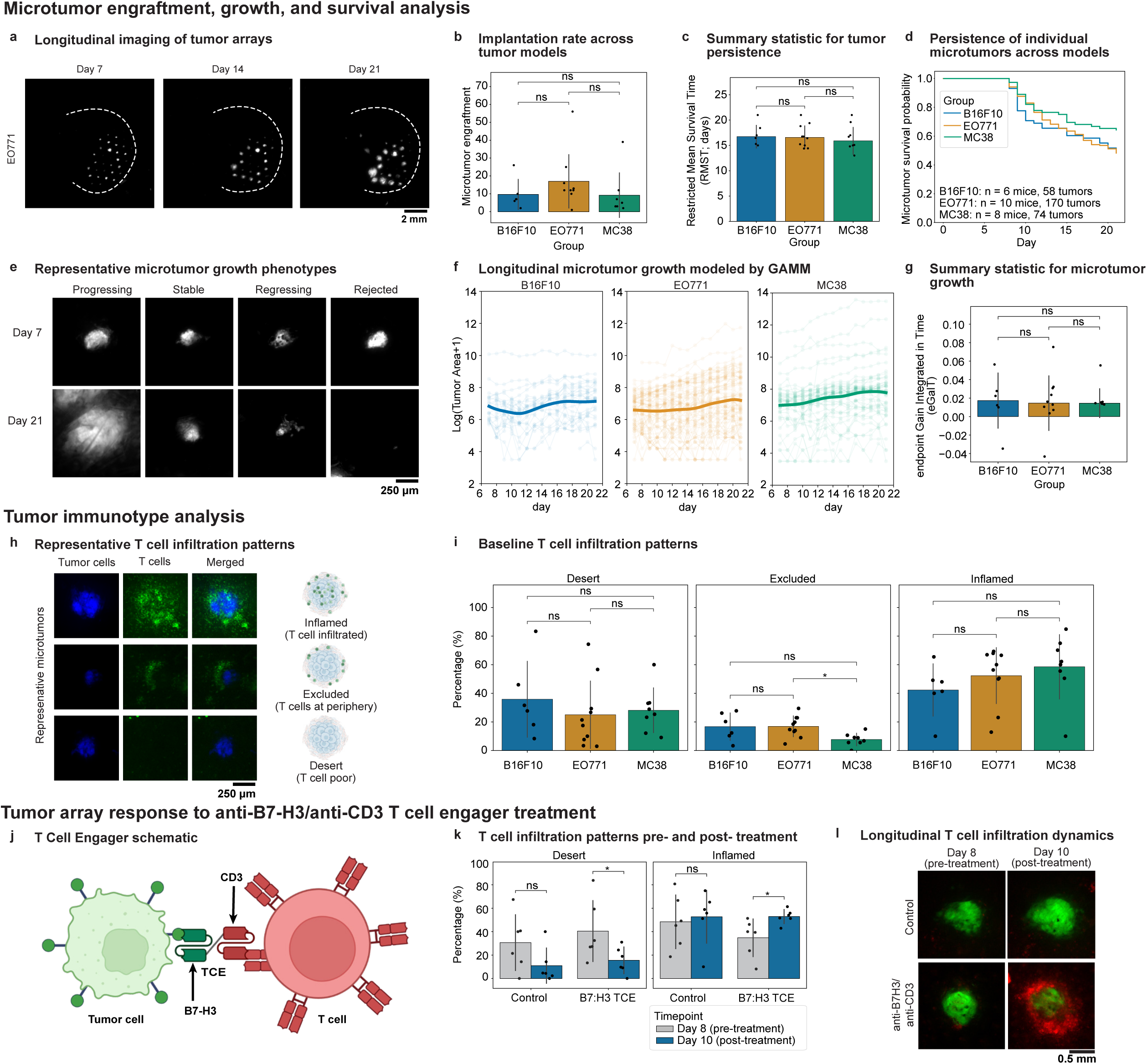
DERM-Array resolves tumor and immune dynamics across models and therapeutic interventions. Tumor arrays of B16F10-mTagBFP2 (n = 6 mice), EO771-mTagBFP2 (n = 10 mice across two independent experiments), or MC38-mTagBFP2 (n = 8 mice) were implanted into Rag2-deficient mice reconstituted with GFP+ CD8+ T cells and analyzed for microtumor growth, persistence, and immune state (**a-i**). **a.** Representative images of an EO771 tumor array on days 7, 14, and 21 post-implantation. Scale bar, 2 mm. **b.** Microtumor engraftment rate per mouse. **c.** Restricted mean survival time (RMST) of microtumors per mouse. **d.** Kaplan–Meier curves showing rejection of individual microtumors, with tumors pooled across all mice within each tumor type. **e.** Representative microtumors exhibiting progressing, stable, regressing, or rejected growth phenotypes on days 7 and 21. Scale bar, 250 µm. **f.** Growth trajectories (log-transformed) of persisting microtumors by tumor type; thin lines represent individual microtumors and bold lines represent the GAMM-fitted summary trajectory for each tumor type. **g.** Endpoint Gain Integrated in Time (eGaIT), a cumulative measure of tumor growth, for persisting microtumors per mouse. **h**. Representative images of tumors (blue) and T cells (green) within an EO771 tumor array exhibiting inflamed, excluded, or desert immunotypes. Scale bar, 250 µm. **i.** Proportions of microtumors assigned desert, excluded, or inflamed immunotype per mouse. **j-l.** Tumor arrays of B16F10-eGFP were implanted into Rag2-deficient mice reconstituted with tdTomato CD8+ T cells and treated with an anti-B7-H3/anti-CD3 T cell engager or PBS control. **j.** Schematic of the bispecific T cell engager (TCE) used for therapeutic intervention, linking B7-H3-expressing tumor cells to CD3+ T cells. **k.** Proportions of TCE treated vs. control microtumors assigned desert or inflamed immunotypes before and after treatment, per mouse. **l.** Representative images of tumors (green) and T cells (red) before (day 8) or after (day 10) treatment. Scale bar, 0.5 mm. For panels **b, c**, **g**, **i**, and **k,** each point represents a single mouse. Bars represent group mean and error bars indicate standard deviation. Statistical analysis was performed using a two-tailed Mann–Whitney U-test.

We next evaluated DERM-Array across multiple tumor models and imaging configurations. B16F10, EO771, and MC38 cells successfully engrafted and exhibited heterogeneous growth and rejection dynamics (**Fig. 2A-G**). DERM-Array also supported imaging of microtumors expressing EGFP, mTagBFP, or VEX, although EGFP-expressing microtumors exhibited higher implantation efficiency and longer persistence than BFP- or VEX-expressing tumors (**Supplementary Fig. 6**). To visualize tumor-immune interactions, we implanted fluorescently labeled tumor cells into Rag2-deficient mice reconstituted with fluorescent T cells. Multicolor imaging resolved distinct patterns of T cell organization, including inflamed, excluded, and desert microenvironments (immunotypes) (**Fig. 2H**). All three tumor models exhibited each immunotype, with differences in their relative frequencies (**Fig. 2I**).

Finally, we used DERM-Array to examine changes in tumor immune organization following immunotherapy. Because B7-H3 has emerged as a promising cancer immunotherapy target (Vitanza et al., 2025), we evaluated an anti-B7H3/anti-CD3 bispecific T cell engager (Zekri et al., 2023; Suita et al., 2026) (**Supplementary Fig. 7**). Systemic administration was associated with changes in the distribution of inflamed, excluded, and desert microtumors (**Fig. 2J**). Longitudinal imaging captured changes in T cell localization within individual lesions following treatment (**Fig. 2K, L**). Together, these experiments demonstrate the ability of DERM-Array to resolve spatial and temporal heterogeneity in tumor immune responses during therapeutic intervention.

An emerging challenge in cancer immunotherapy is the heterogeneity of treatment responses within individual patients. Separate metastatic tumors can respond differently to the same systemic treatment (interlesional heterogeneity; Reuben et al., 2017), while distinct regions within a single tumor can exhibit different patterns of immune infiltration and function (intratumoral heterogeneity; AbdulJabbar et al., 2020; Nirmal et al., 2022). Recent preclinical models have begun to uncover the mechanisms underlying this variability by studying distinct immune microenvironments within a single host. Li et al. (2018) demonstrated that differences in tumor cell signaling can establish immune inflamed or noninflamed microenvironments and identified CXCL1-mediated myeloid recruitment as a mechanism of immune exclusion. Dhainaut et al. (2022) extended this approach through spatial genetic screening to systematically identify tumor genes that regulate local immune and stromal organization. Ortiz-Muñoz et al. (2023) then introduced STAMP and demonstrated that distinct immune microenvironments and tumor outcomes can emerge even among neighboring tumors initiated from the same cell line, identifying additional local cellular interactions that promote T cell infiltration and response to immunotherapy. Most recently, Tanaka et al. (2025) demonstrated that immune-cold tumor-cell subpopulations can establish localized immunosuppressive microenvironments that persist following immunotherapy and serve as nucleate sites for treatment resistance. Together, these studies reveal that spatial immune heterogeneity can arise through multiple mechanisms, identify potential therapeutic targets for overcoming local immune suppression, and highlight the need to develop treatments that remain effective across diverse tumor immune microenvironments.

Tumor arrays are particularly well suited to this challenge because they allow individual tumors to be imaged repeatedly within a shared host environment, linking early immune organization to subsequent tumor growth, rejection, and therapeutic response. Building on the original STAMP platform (Ortiz-Muñoz et al., 2023; Carbone et al., 2026), DERM-Array makes this experimental approach accessible to the broader research community through commercially available hardware and open source protocols. The method can be readily adapted to different tumor models and treatment strategies, allowing investigators to address the specific mechanisms of immune resistance relevant to their disease and therapeutic context. We envision tumor arrays becoming a widely adopted platform for understanding how spatial immune heterogeneity emerges and evolves and for developing treatments that achieve effective tumor control across diverse immune microenvironments.

## Methods

### Animals and reagents

#### Mice

Animals were housed in specific pathogen-free conditions at the Seattle Children’s Research Institute (SCRI) Building CURE Animal Facility. SCRI is an AAALAC-approved institution, and all procedures were performed according to protocols approved by Institutional Animal Care and Use Committee (IACUC). Mice were socially housed (maximum five same sex adults per cage, except when breeding) in NexGen™ Mouse 500 AllenTown caging systems; cages were individually vented and maintained on an automatic water system with chlorinated water. Cages were supplied with corn cob bedding with Nestlet™ and crinkle paper as enrichment; a standard diet of PMI PicoLab Rodent 5053 was provided via food hopper. Cage changes occurred every 2 weeks, with spot changes as needed. Housing rooms were maintained at a temperature of 68-79°F (20.0 to 26.1 °C), 30-70% relative humidity, and a lighting schedule of 12 hours on, 12 hours off. Mice were identified by paw pad tattoos rather than ear tags or punches. Tattoo ink (Amazon ASIN B07HQPPZY6) was applied using a 27G needle while mice were anesthetized or restrained. Individual mice were identified by combinations of tattooed paws. Healthy adult mice aged six to fifteen weeks, with no apparent abnormalities were used for all experiments. Animal sex was selected according to tumor model (B16F10 and MC38, male; EO771, female), unless otherwise indicated.

Wild-type C57BL/6J mice (JAX #000664) were purchased directly from JAX for tumor growth and survival experiments. For immune phenotyping experiments, C57BL/6J-Rag2em3Lutzy/J (Rag2–/–; JAX #033526) mice were used as recipients for adoptive T cell transfer. T cell donors were C57BL/6-Tg(CAG-EGFP)131Osb/LeySopJ (JAX #006567) mice or mice with constitutive tdTomato expression generated by crossing B6.Cg-Gt(ROSA)26Sortm14(CAG-tdTomato)Hze/J (JAX #007914) mice with CMV-Cre mice (JAX #006054) in house. Breeding colonies of Rag2–/– recipients and both fluorescent T cell donor lines were maintained in-house to generate experimental cohorts.

#### Cell lines

B16F10 melanoma cells (CRL-6475; obtained from ATCC) and MC-38 colon carcinoma cells (T8291; obtained from ABM) were cultured in media consisting of Dulbecco’s Modified Eagle’s Medium (DMEM; Thermo Fisher Scientific #11960044), 1% GlutaMAX (100X; Thermo Fisher Scientific #35050-061), 1% penicillin-streptomycin (P/S; Thermo Fisher Scientific #15070063) (100X), and 10% FBS (Fisher Scientific #A5256801). EO771 mammary carcinoma cells (T1302; obtained from ABM) were cultured in the media described with the addition of 1% HEPES (1M; Thermo Fisher Scientific #15630080) and 4% 7.5% sodium bicarbonate (Thermo Fisher Scientific #25080094).

All cell lines were engineered to express fluorescent proteins eGFP, mTagBFP2, and VEX using the piggyBac transposase (System Biosciences #PB210PA-1) system delivered via lipofection (Lipofectamine 3000; ThermoFisher #L3000001) according to manufacturer recommendations. Cargo plasmids were synthesized, propagated, and assessed for quality by GenScript. Fluorescently labeled cells were purified on a BD FACSymphony S6 cell sorter.

Each engineered cell line was optimized for in vivo growth via *in vivo* passage. Tumor cells were implanted in ear tissue via laser ablation and/or flank via subcutaneous injection and harvested between days 9-21. Tumors were harvested and enzymatically dissociated in RPMI 1640 containing collagenase P (0.2 mg/mL; Roche, Cat. No. 11213865001) and DNase I (0.1 mg/mL; Millipore Sigma, Cat. No. 10104159001) for 15 min at room temperature, followed by mechanical dissociation and resuspension in the appropriate cell culture medium. Recovered tumor cells were cultured as described above. After at least one week in culture, a fluorophore-positive cell population was purified on a BD FACSymphony S6 cell sorter.

### DERM-Array methodological development and validation

#### Ex vivo laser ablation

Mouse ears were depilated using Veet hair removal cream (Amazon B001KYPZ4G), excised, and fixed in 4% paraformaldehyde (Electron Microscopy Sciences #15712-S) for 12-24 h at 4 °C. Tissues were then stored in phosphate-buffered saline or 70% ethanol at 4 °C for up to one month before laser treatment. Immediately prior to laser ablation, ear tissues were thoroughly dried, then mounted on acrylic trays with the ventral side facing upwards using double-sided tape (Tesa 4970). A thin layer of charcoal powder (Millipore Sigma #242276-250G) was applied to each tissue with a latex-free wedge-style cosmetic sponge (Amazon B0BNZZQZN2).

Prepared tissues were placed in the xTool F1 Ultra (Makeblock Co., P5010341) workspace, and the array file was processed with user-defined settings:

Object style: Engrave
Laser type: Blue light
Power (%): 20-60
Speed (mm/s): 10
Pass: 1
Lines per cm: 300
Engraving mode: Uni-directional
Engraving angle: Incremental; 0
Cross hatch: Disabled

#### FITC-dextran labeling and mounting

Excised mouse ears were prepared and subjected to laser ablation as described above, excluding PFA-fixation. Tissues were incubated with 100 µL of 1 mg/ml FITC-dextran (Millipore Sigma #FD4-100MG) in PBS for two hours at 4°C, then vigorously washed with 70% ethanol, then PBS. Custom imaging chambers were assembled using glass microscope slides (Thermo Fisher Scientific, #12-550-A3) and gaskets prepared from adhesive silicone sheets (Amazon, ASIN B0B4X47N19). Double-sided tape (Tesa 4970) was applied to the non-adhesive side of the silicone sheets, and individual gaskets with 20 mm circular openings were cut using a leather hole punch (Amazon B0FLQ19L61). The gaskets were attached to microscope slides, and ear tissues were secured to the glass within each opening using cyanoacrylate adhesive (Loctite 12564) with the laser ablated surface facing upward. Once dry, 200 µl PBS was added to cover the tissue and a No. 1.5 coverslip (Ibidi 10812) was adhered to the upper gasket surface to seal the chamber.

#### Confocal depth measurement

Confocal z-stacks of individual implantation sites were acquired using a Zeiss LSM900 confocal laser scanning microscope (FITC, 10x objective). Implantation site depth was determined from the axial distance between the surrounding, non-ablated skin surface and the base of each implantation site, identified by FITC-dextran fluorescence. Depth was calculated from the number of z-steps separating these positions and the acquisition step size (Fiji 2.16.0/1.54t).

#### Implantation site area measurement

Implantation site arrays were imaged on the Leica M205 FA stereomicroscope in brightfield configuration and through a Cy3 filter under ambient light to capture autofluorescence. Images were analyzed with Fiji; implantation site interiors were segmented with a custom labkit classifier followed by manual curation and implantation site areas were quantified using native Fiji (2.16.0/1.54t) measurement tools.

#### Histology

Excised mouse ears were prepared and subjected to laser ablation as described above. Tissues were stored in 70% ethanol before paraffin embedding. Ears were oriented perpendicular to the block surface to obtain transverse cross sections through the implantation sites. Paraffin blocks were sectioned at 5 µm and sections containing implantation sites were stained with hematoxylin and eosin.

#### Histological image quantification

H&E-stained sections were imaged on the Evident VS200 (brightfield, 20x). The depth and diameters of individual implantation sites were manually annotated and quantified using QuPath (0.7.0).

### In vivo tumor array experiments

#### DERM-Array laser ablation

One to three days prior to laser ablation, mice were anesthetized with isoflurane and hair was removed from both surfaces of ear pinnae using Veet hair removal cream (Amazon B001KYPZ4G). After 1-3 minutes, the cream was removed with a Kimwipe and residual product was rinsed off with warm water.

On the day of laser ablation, mice were anesthetized by intraperitoneal injection of ketamine (100 mg/kg; Patterson #07-894-8462) and xylazine (15 mg/kg; Patterson #07-891-6165) in sterile saline. Mice were placed on a heating pad (37°C) to maintain body temperature and ophthalmic ointment (Systane #0065050939) was applied to prevent corneal desiccation. Mice were positioned supine on custom trays constructed from 1/2-inch-thick black acrylic sheets (8×8 inches; TAP Plastics). Strips of double-sided adhesive tape (Tesa 4970) were placed on the trays to secure the head and ears. Both ears were extended away from the head and adhered to the tape with their dorsal surfaces facing down, leaving the ventral surfaces exposed for laser ablation. A thin layer of charcoal powder (MilliporeSigma #242276-250G) was applied to the exposed ear ventral surfaces using a latex-free cosmetic sponge (Amazon B0BNZZQZN2). Mice were then transferred to the F1 Ultra (xTool) laser workspace for ablation.

The xTool Creative Studio application was used to position array vectors and initiate laser ablation. The array file was processed with user-defined settings:

Object style: Engrave
Laser type: Blue light
(%): 30-40%
Power (mm/s): 10
Speed Pass: 1
per cm: 300
Engraving mode: Uni-directional
Engraving angle: Incremental; 0
Cross hatch: Disabled

For the first three days after laser ablation, petroleum jelly (Medline CUR005345) was applied daily to the ears using a cotton swab to prevent desiccation and scabbing.

#### STAMP laser ablation

For STAMP experiments, mice were depilated and anesthetized as described above excluding charcoal powder application. Anesthetized mice were maintained on a heating pad and positioned on rectangular plastic trays (Amazon B0CP2BK67J) as described above with strips of double-sided adhesive tape securing the dorsal surface of the ears flat against the tray. Implantation sites were generated on the ventral ear surface using a P.L.E.A.S.E. laser system (Pantec Biosolutions), as previously described (Ortiz-Muñoz et al., 2023; Carbone et al., 2026). The laser handpiece was manually positioned perpendicular to the ear surface and triggered separately for each ear. The laser was operated using the following custom STAMP program settings:

Target ablation depth: 91 µm
Pulses per implantation site: 2
Power: 0.9 W
Frequency: 200 Hz
Pulse duration: 125 µs
Density: 9%
Array size: 14 × 14

The device filter was replaced as needed to maintain laser efficiency above 75%.

#### Tumor cell preparation and implantation

Tumor cells were cultured in tissue culture-treated flasks (Stellar Scientific NST-721003) until approximately 80-100% confluent. To prepare for implantation, cells were detached with 0.25% Trypsin (Thermo Fisher Scientific 25200056), centrifuged at 500 × g for five minutes, resuspended in sterile phosphate-buffered saline (Corning 46-013-CM), and kept on ice. Immediately after laser ablation, tumor arrays were seeded by pipetting 100 µL cell suspension containing 2.5-10.0×10^6^ cells onto each ear. After a thirty-minute incubation, excess cell suspension was removed with a Kimwipe and mice were returned to their home cage.

#### T cell isolation and transfer

T cells were isolated from spleens of eGFP or tdTomato-expressing donor mice. CD8+ T cells were isolated with the EasySep Mouse CD8+ T cell Isolation Kit (Stem Cell #19853) according to the manufacturer’s instructions. T cell transfers were performed with a target dose of 0.7-1.5×10ª purified CD8+ T cells per recipient, using approximately one donor spleen for every three to four recipients. Cells were administered by retro-orbital injection at least one day before tumor implantation.

#### In vivo fluorescence imaging

In vivo tumor array imaging was performed on a Leica Microsystems M205 FA fluorescence stereomicroscope (10450040) equipped with a PlanApo 1.0x objective (Leica 10450028), 10x eyepiece (10450910), Leica XY-Scanning stage (10450665), and a Leica K8 camera (11547116). Fluorescence imaging was conducted using ET BFP (mTagBFP2; Leica 10450329), ET Fluorescein (eGFP and VEX; Leica 10450646), and ET Cy3 (tdTomato; Leica 10450198) filter sets. VEX tumor arrays were illuminated using ultraviolet flashlights (394 nm; Amazon B0B92PDCSV) mounted on the gooseneck filters and emissions were captured through the ET Fluorescein filter. Images were acquired and analyzed using LAS X Premium software (Leica 11640717).

Mice were anesthetized with inhalant isoflurane delivered via induction chamber, then transferred to a nose cone attached to the microscope stage for continued immobilization. Double sided tape (Thomas Scientific #1520CL100P3D) was used to secure the dorsal side of mouse ears to the microscope stage to expose the ventral surface for imaging. A total magnification of 10x was used for tumor-only experiments, whereas immune phenotyping experiments involving fluorescent T cells were imaged at 30x magnification.

### Therapeutic intervention studies

#### TCE design and production

Single-chain variable fragments (scFvs) derived from the anti-murine B7-H3 antibody clone m276 (Haydar et al., 2021) and the anti-murine CD3 antibody clone 2C11 (Leo et al., 1987) were joined by a short Gly-Ser (GS) linker to generate a tandem scFv B7-H3xCD3 T-cell engager (TCE) with a C-terminal hexa-histidine (6XHis) tag. The TCE was produced using the Daedalus mammalian expression system (Bandaranayake et al., 2011). The coding sequence was codon-optimized for human expression, synthesized, and cloned into the Daedalus mammalian expression vector VCR020 (GenScript). A second-generation lentiviral packaging system was used to generate lentivirus encoding the TCE, which was then used to transduce HEK293F cells. Secreted TCE was recovered from the culture supernatant by affinity chromatography using a HisTrap FF crude column (Cytiva, catalog no. 17-5286-01), followed by preparative size-exclusion chromatography using a HiLoad 26/600 Superdex 200 pg column (Cytiva, catalog no. 28-9893-36) on an ÄKTA FPLC system. The final formulation buffer was 1X phosphate-buffered saline (PBS), pH 7.4. Purified material was characterized by analytical size-exclusion chromatography using a Superdex 200 Increase 10/300 GL column (Cytiva, catalog no. 28-9909-44) and by SDS-PAGE under reducing and nonreducing conditions. Final protein purity was >95%. Endotoxin was measured with an Endosafe nexgen-PTS system (Charles River) and was below 0.5 EU mg̅¹ protein.

#### In vitro T cell engager killing assay

For in vitro cytotoxicity assays, B16F10, MC38, and EO771 expressing eGFP, as previously described, were used as target cell lines. Cells were plated at 2.0×10^4^ cells per well in 96-well plates (Nunc Edge 167425) and rested for 24 hours. Murine primary splenic T cells isolated from female C57BL/6 mice were expanded with anti-mouse CD3ε antibody (clone 145-2C11, Biolegend 100339, 5ug/ml) and IL-2 (Peprotech 200-02, 10ng/ml) for 7 days then co-cultured with target cells at a 5:1 effector:target ratio. The anti-B7-H3/anti-CD3 T cell engager was added across a five-fold dilution series from 100 nM to 1.28pM in triplicate. GFP-positive object counts were acquired over 48 hours with an Incucyte SX5 Live-Cell Analysis System (Sartorius). Specific lysis was calculated relative to baseline and normalized to effector-plus-target control lacking bispecific. Values were determined via fitting a three-parameter sigmoid to the 48-hour data in GraphPad Prism (version 11.0.2, GraphPad Software).

#### T cell engager treatment

Male Rag2–/– mice received tdTomato-expressing CD8+ T cells and B16F10-eGFP tumor arrays as described above. Mice were assigned to treatment or control groups based on initial tumor burden, with consideration of cage effects. Beginning on day 8 after tumor implantation, mice were imaged daily to assess tumor growth and T cell spatial organization. On day 9, mice received a single intraperitoneal injection of 1 µg anti-B7-H3/anti-CD3 bispecific T cell engager or PBS. Imaging continued through day 10.

### Transcriptomic profiling

#### Sample preparation

Animals were subjected to laser ablation only (day 7, day 3, day 1, n = 1 ear), laser ablation with tumor cell seeding (tumors day 14, n = 2 ears; tumors day 7, day 3, n = 1 ear), or no intervention (healthy; n = 1 ear) on one or both ears. Four-millimeter biopsy punches (Electron Microscopy Sciences #69031-05) of middle-outer ears were collected immediately after animal euthanasia, placed into bead mill homogenizer tubes (Revvity #19-628) filled with 1 ml cold TRIzol Reagent (Thermo Fisher Scientific #15596026), and kept on wet ice. Homogenization was performed using Omni Bead Ruptor Elite (Revvity #19-042E); optimal homogenization was achieved with one 30 seconds of homogenization at 7.45 m s-1, 1-3 minutes cooling on wet ice until cold to the touch, then an additional round of 15 seconds at the same speed. Homogenized samples were centrifuged at 12,000 x g for 5 minutes, then lysate was transferred to a 1.5 ml Eppendorf and frozen at -80°C overnight.

Briefly, bulk RNA extraction included phase separation (chloroform: Fisher Scientific #AAJ67241AP) and isopropanol (Fisher Scientific #AC327272500) precipitation with ethanol wash (MilliporeSigma #1085430250) according to manufacturer instructions. RNA was purified using the GeneJET RNA Purification Kit (Thermo Fisher Scientific #K0842) with on-column DNase I digest (Thermo Fisher Scientific #EN0521) according to manufacturer instructions. To assess quantity and quality of RNA, we used the Qubit RNA HS (Thermo Fisher Scientific #Q32852) and IQ Assay kits (Thermo Fisher Scientific #Q33221) measured with a Qubit 4.0 Fluorometer (Thermo Fisher Scientific #Q33226). Samples in molecular biology grade water were prepared for submission via room-temperature drop box by adding 10X TE buffer (Invitrogen #T11493) and SEQguard™ Dino Preserve (Genovis) as directed, then bulk RNA sequencing was performed by PlasmidSaurus.

#### Bulk RNA-seq analysis

Principal component analysis (PCA) was performed on log2(CPM + 1) transformed expression values from the 2,000 most variable protein-coding genes, filtered for mean expression ≥1 CPM and expression ≥1 CPM in at least three samples.

To characterize the temporal progression of tissue injury, inflammation, and immune activation following laser ablation, custom gene expression signatures were constructed from mouse GO Biological Process 2026 and MSigDB Hallmark 2020 gene sets obtained through Enrichr. The three measured signatures were constructed by pooling functionally related gene sets reflecting top differentially expressed genes in our dataset: (1) skin injury response (GO:0042060, GO:0090594, GO:0030216); (2) neutrophil/IL-1 inflammation (GO:0030593, GO:0042119, GO:0032731, GO:0071347); and (3) interferon response (Hallmark interferon-α and interferon-γ responses). After removing duplicates and filtering expression (mean ≥1 CPM and ≥1 CPM in at least three samples), the 15 genes with the greatest maximum log2 fold change relative to healthy skin across post-ablation timepoints were selected for each signature. Expression values were log2(CPM + 1) transformed and standardized to gene-wise z-scores across samples. Summary signature scores were calculated as the mean z-score of the 15 selected genes per sample, while heatmaps display the individual gene-wise z-scored expression values of the same 15 genes comprising each signature.

### Computational image analysis

#### Microtumor segmentation model training

A two-dimensional nnU-Net v2 model was trained for automated microtumor segmentation using 2,394 paired fluorescence images and manually annotated binary segmentation masks collected across 17 independent experiments. Training data were prepared using custom Python scripts in Jupyter notebooks, and model training was performed on the full dataset using the nnU-Net v2 framework (Isensee et al., 2021). The trained model was applied to new fluorescence imaging datasets using custom Python scripts to generate initial tumor segmentation masks, which were subsequently manually reviewed and curated using Labkit (v0.4.0) in Fiji (v2.16.0/1.54t).

#### Tumor survival and growth analysis

Survival analysis included restricted mean survival time (RMST; restricted_mean_survival_time from lifelines 0.30.0) calculations for all tumors within each mouse, then group-level differences were assessed using two-tailed Mann-Whitney U-tests. The Cox Proportional Hazards model (CPHFitter from lifelines, 0.30.0) was used to compare pooled microtumor persistence across groups, with robust variance estimation and clustering by mouse ID.

To compare microtumor growth, we employed the strategy described in Forrest et al. (2020). Briefly, generalized additive mixed models (GAMM) were fitted to pooled growth trajectories of persisting tumors using pyGAM (GAM, 0.12.0). Endpoint Gain Integrated in Time (eGaIT) values were calculated for all persisting tumors per mouse, then group-level differences were assessed using two-tailed Mann-Whitney U-tests.

See Supplemental Note 1 for a detailed discussion of microtumor growth and survival analysis.

#### T cell infiltration analysis

T cell infiltration patterns (“immunotypes”) were quantified using median radial fluorescence profiles generated from tumor regions of interest, as previously described (Ortiz-Muñoz et al., 2023; Carbone et al., 2026). Microtumors were classified as desert, excluded, or inflamed using custom Python scripts in Jupyter notebooks. Tumors with median T cell fluorescence below an intensity threshold of 200 were classified as desert. For the remaining tumors, radial fluorescence profiles were min-max normalized, and the ratio of median fluorescence in the inner 25% of the radial profile to that in the remaining peripheral region was calculated. Tumors with a core-to-periphery fluorescence ratio below 0.8 were classified as excluded, while those with ratios ≥0.8 were classified as inflamed.

### Statistics and reporting

#### Statistical analysis and reproducibility

Mice from different treatment groups or tumor cell line cohorts were housed together to minimize cage effects. For T cell engager (TCE) experiments, mice were assigned to treatment and control groups to balance pretreatment tumor burden. Investigators were blinded to experimental groups during tumor segmentation.

Statistical analysis was performed using two-tailed Mann-Whitney U-tests (statsannotations 0.7.2). p < 0.05 was significant. Bonferroni correction was applied when performing multiple comparisons. Unless otherwise indicated, n refers to individual mice.

See Supplemental Note 1 for a detailed discussion of statistical methods.

## Figure Preparation

All figures were assembled in Adobe Illustrator 2026. Figs. 1a, 2h, and 2j were created using BioRender. Data was prepared for plotting using numpy (1.26.4), pandas (2.3.3), and scipy (1.16.3). Plots were generated in Jupyter Lab (4.5.2) using Python packages including MatPlotLib (3.10.8), Seaborn (0.13.2), and previously mentioned packages. Some plots were prepared using GraphPad Prism (11.0.2).

## Code availability

Custom code and analysis workflows used in this study will be made publicly available through an open-access repository upon publication.

## Data availability

The data supporting the findings of this study, including source data and sequencing datasets, will be deposited in appropriate public repositories and made available upon publication. Repository accession numbers and links will be provided in the final manuscript.

## Acknowledgements

We thank M. Coleman for tissue supply; T. Greywal, J. Tsai, and members of the R. Ram lab for methodological discussions; A. Chithran for confocal imaging guidance; G. Schultz for histology; the Seattle Children’s Research Institute Office of Animal Care and Veterinary team for animal care.

## Ethics declarations

The authors declare no competing interests.

## Author contributions

Imaging: M.M., S.H., K.F., R.D., A.M., J.M.; Coding: M.M., S.H.; Data Analysis: M.M.; Conceptualization and design: M.M., J.P.P, C.B.C.; Animal husbandry: S.H.; Reagents: K.F., J.P.P., R.R.; TCK assay: R.R., S.S.

## Supplementary Information

**Supplementary Figure 1.**
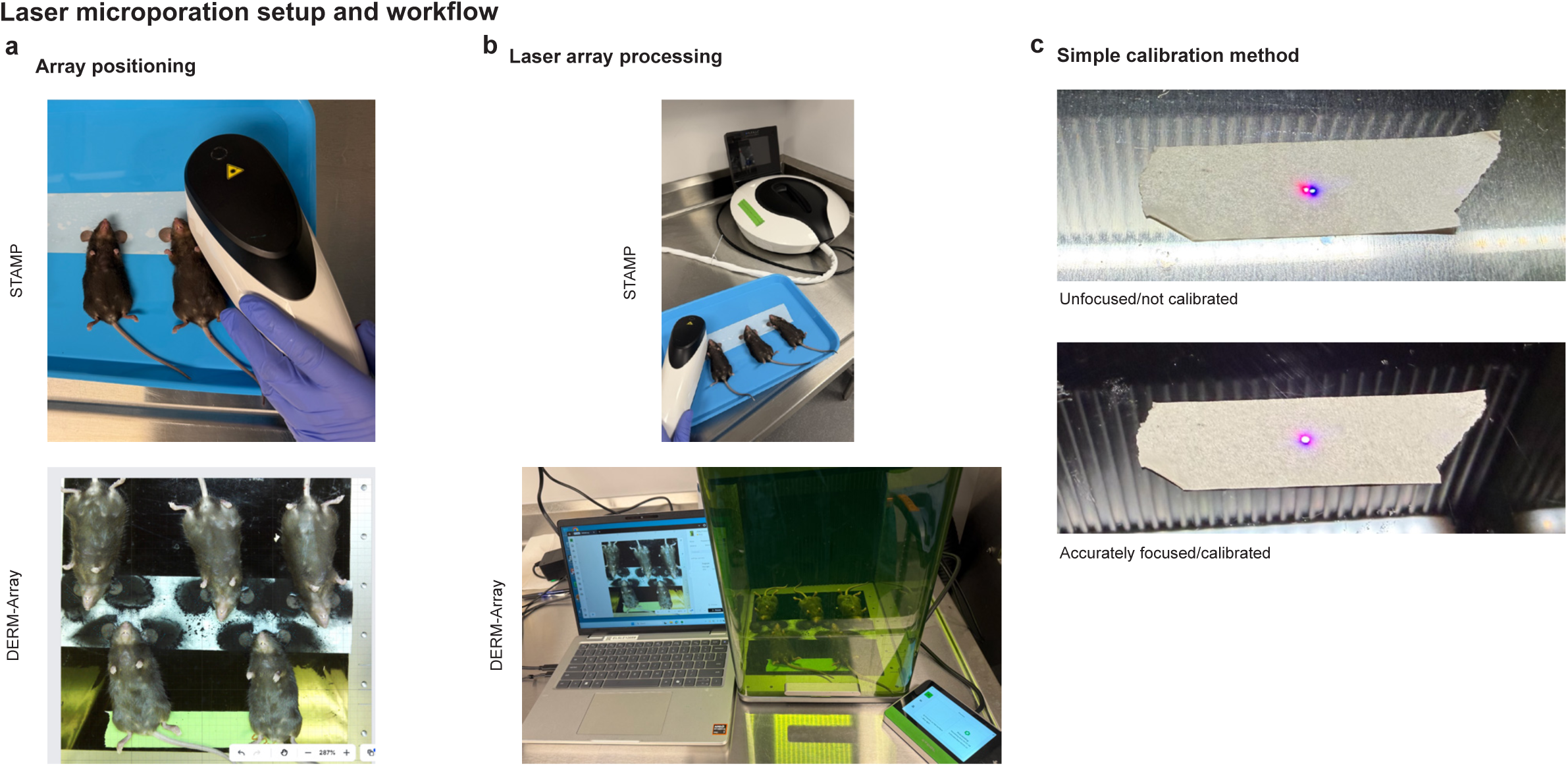
DERM-Array streamlines the laser ablation workflow. **a.** Array positioning method using the STAMP handpiece and the DERM-Array software interface. **b.** Experimental setup during laser ablation. **c.** DERM-Array laser device calibration using red and blue laser alignment markers projected onto white tape. Overlap of red and blue markers indicates calibration accuracy.

**Supplementary Figure 2.**
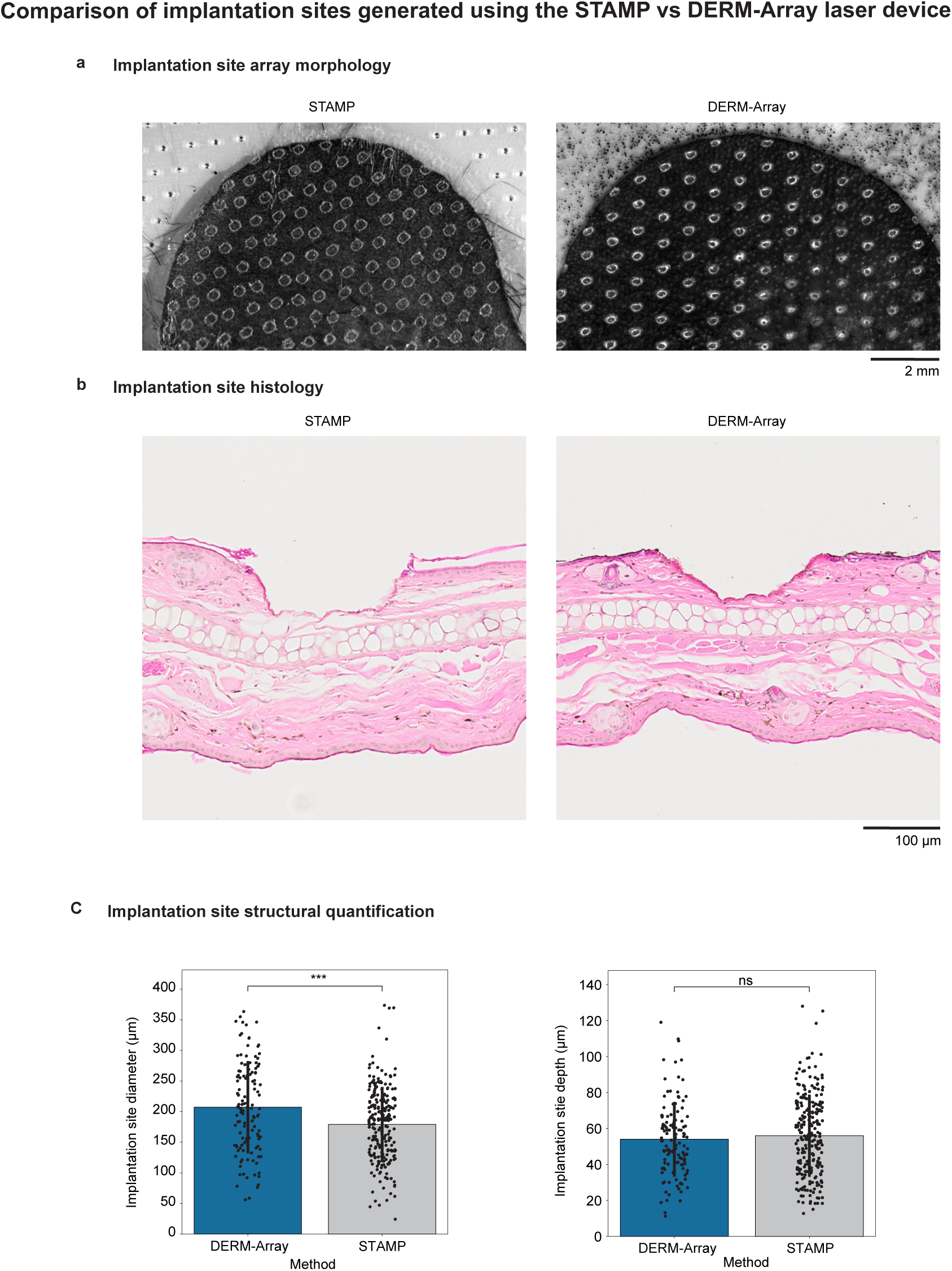
DERM-Array recapitulates STAMP implantation site architecture using an inexpensive diode laser. **a.** Representative images of implantation site arrays generated in excised ear pinnae tissue using STAMP or DERM-Array. **b.** H&E-stained cross-section of an implantation site in FFPE tissue. **c.** Quantification of implantation site diameter and depth measured from H&E images. Each point represents a single implantation site (STAMP, n = 240 sites from 5 ears; DERM-Array, n = 125 sites from 5 ears). Bars represent group mean and error bars indicate standard deviation. Statistical analysis was performed using a two-tailed Mann–Whitney U-test.

**Supplementary Figure 3.**
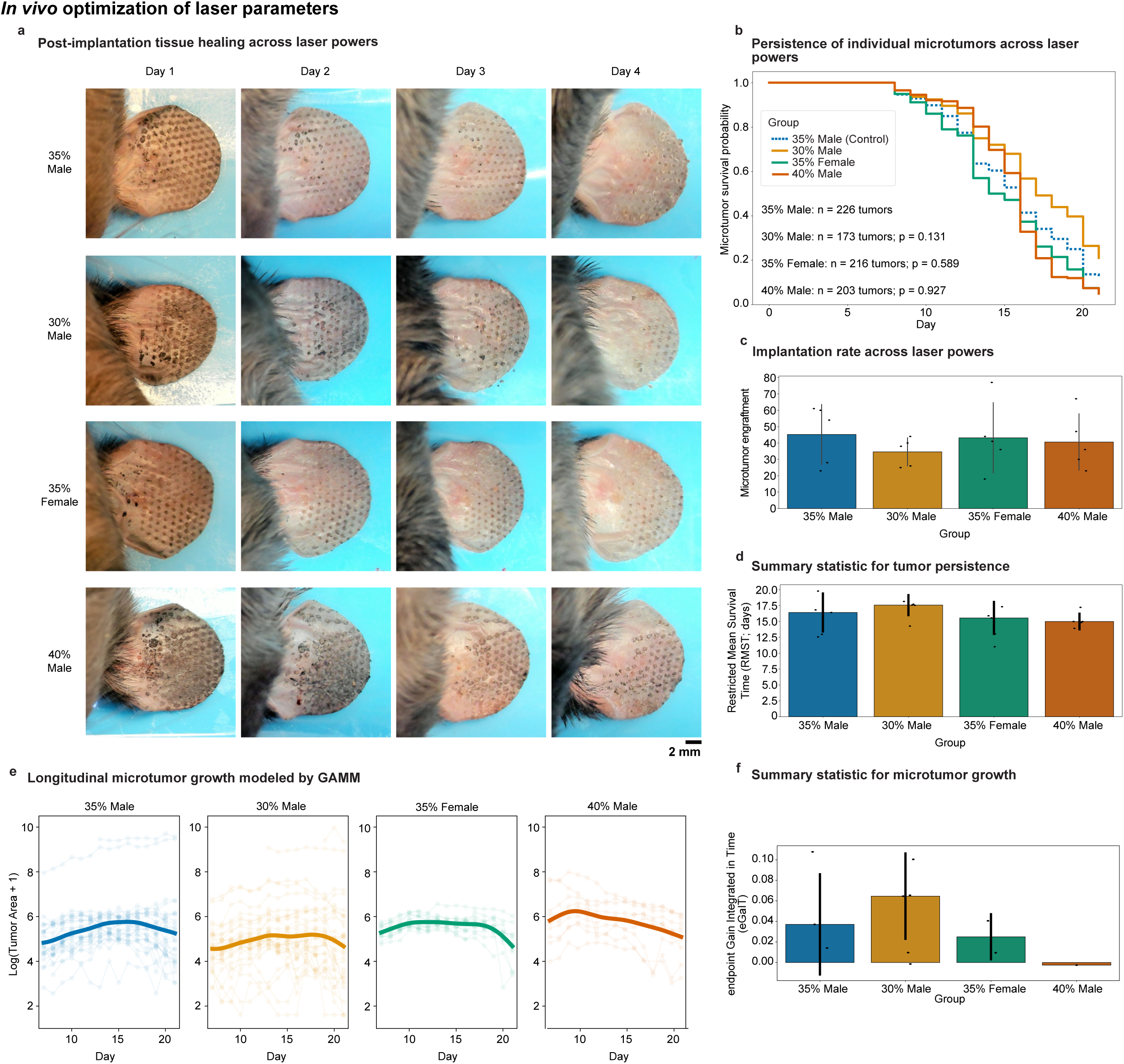
DERM-Array supports tumor array generation across a practical range of laser parameters. **a.** Representative images of ear pinnae showing tissue healing for four days post-procedure across laser powers (30-40%; 6-8 Watts). n = 5 mice per laser power. Scale bar, 2 mm. **b.** Kaplan–Meier curves showing rejection of individual microtumors, with tumors pooled across all mice within each laser power. **c.** Microtumor engraftment rate per mouse. **d.** Restricted mean survival time (RMST) of microtumors per mouse. **e.** Growth trajectories (log-transformed) of persisting microtumors by laser power; thin lines represent individual microtumors and bold lines represent the GAMM-fitted summary trajectory for each laser power. **f.** Endpoint Gain Integrated in Time (eGaIT), a cumulative measure of tumor growth, for persisting microtumors per mouse. For panels **c**, **d,** and **f,** each point represents a single mouse. Bars represent group mean and error bars indicate standard deviation. Statistical comparisons for engraftment, RMST, and eGaIT were performed using a two-tailed Mann– Whitney U-test; hazard ratios and p-values for survival comparisons were calculated using Cox Proportional Hazards Model.

**Supplementary Figure 4.**
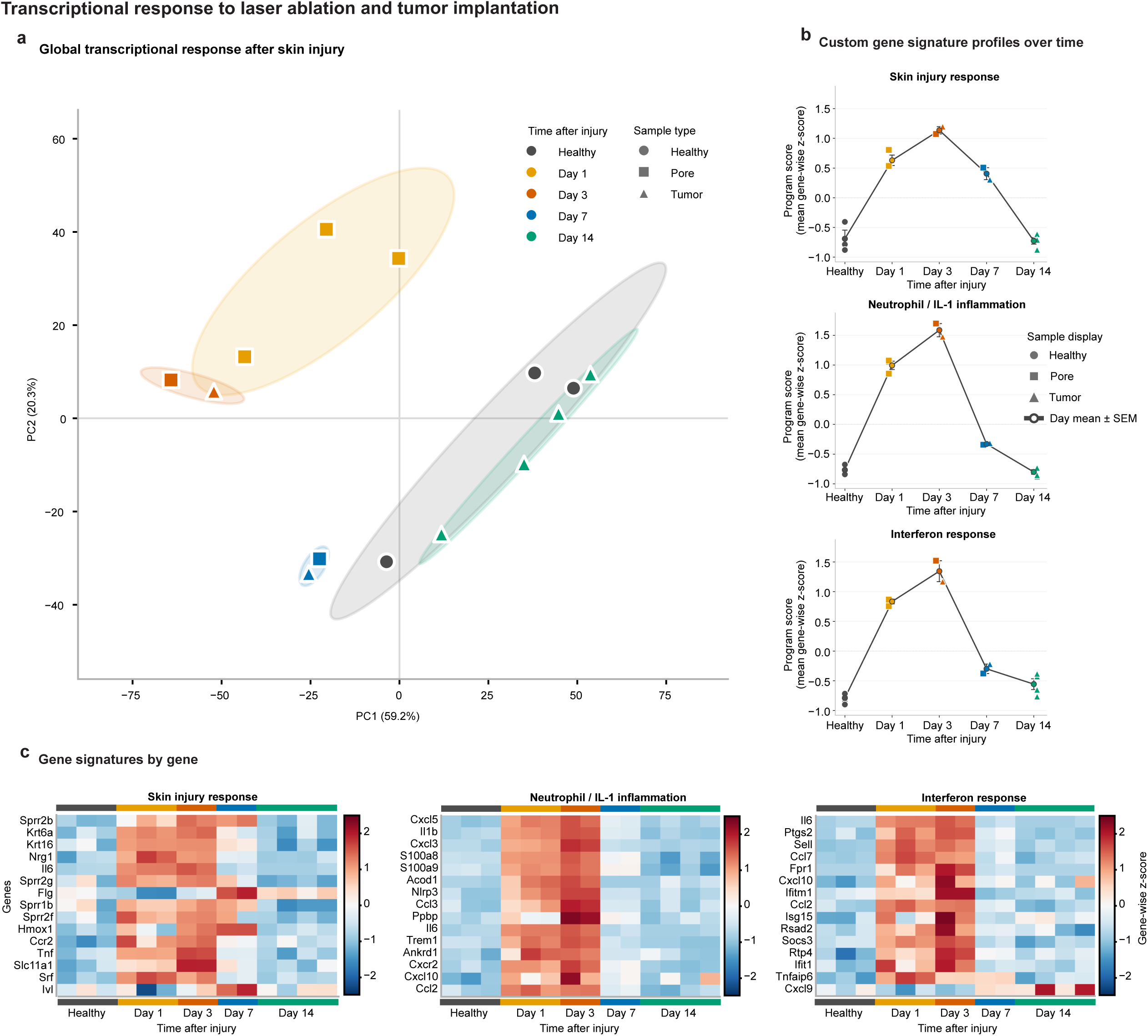
Inflammatory tissue response to laser ablation normalizes over time. **a-c.** WT mice received laser ablation with or without tumor implantation. Tissue biopsy punches were collected on days 1, 3, 7, or 14 and analyzed by bulk RNA sequencing. Gene expression was compared to healthy skin controls. **a.** Principal component analysis (PCA) of transcriptomic profiles across timepoints. **b.** Temporal changes in gene expression signatures associated with skin injury, neutrophil/IL-1 inflammation, and interferon responses, calculated as the mean gene-wise z-score of log2(CPM + 1) expression across the genes comprising each signature. Points represent individual samples, lines connect timepoint means, and error bars indicate SEM. **c.** Heatmap showing gene-wise z-scored expression of the 15 genes comprising each signature in **b** across timepoints. Skin injury response (left), neutrophil/IL-1 inflammation (middle), and interferon response (right).

**Supplementary Figure 5.**
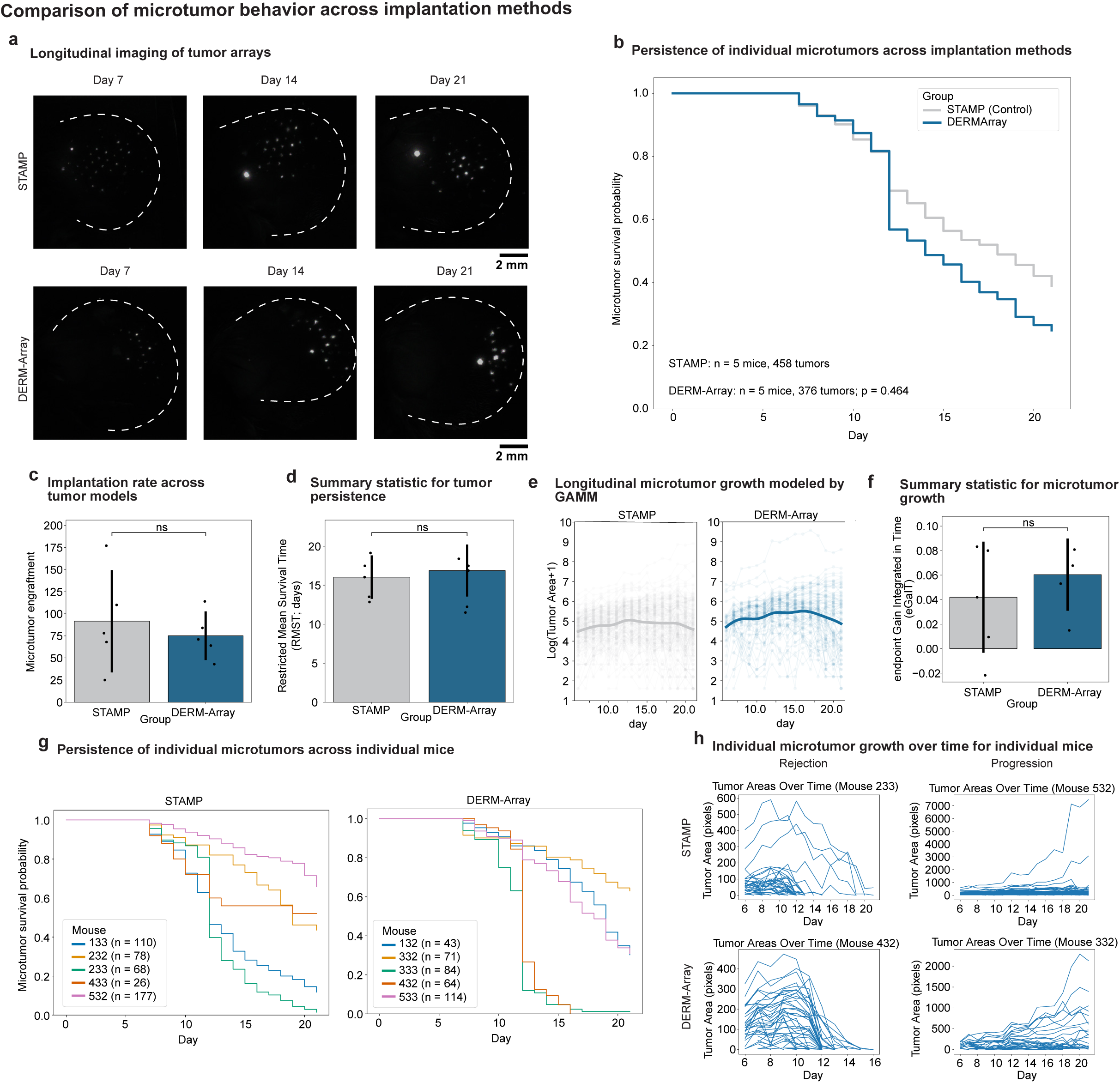
Tumor behavior dynamics are conserved across implantation methods. B16F10-eGFP tumor arrays were implanted in WT mice using STAMP or DERM-Array (n = 5 mice per group). **a.** Representative images of tumor arrays on days 7, 14, and 21 post-implantation. Scale bar, 2 mm. **b.** Kaplan–Meier curves showing rejection of individual microtumors, with tumors pooled across all mice within each implantation method. **c.** Microtumor engraftment rate per mouse. **d.** Restricted mean survival time (RMST) of microtumors per mouse. **e.** Growth trajectories (log-transformed) of persisting microtumors by tumor type; thin lines represent individual microtumors and bold lines represent the GAMM-fitted summary trajectory for each tumor type. **f.** Endpoint Gain Integrated in Time (eGaIT), a cumulative measure of tumor growth, for persisting microtumors per mouse. **g.** Kaplan–Meier curves showing rejection of all individual microtumors pooled per mouse. **h.** Representative microtumor growth trajectories from individual mouse ears exhibiting predominantly rejecting or predominantly progressing tumor arrays. For panels **c**, **d**, and **f**, each point represents a single mouse. Bars represent group mean and error bars indicate standard deviation. Statistical analysis was performed using a two-tailed Mann–Whitney U-test; hazard ratios were calculated using Cox Proportional Hazards Model.

**Supplementary Figure 6.**
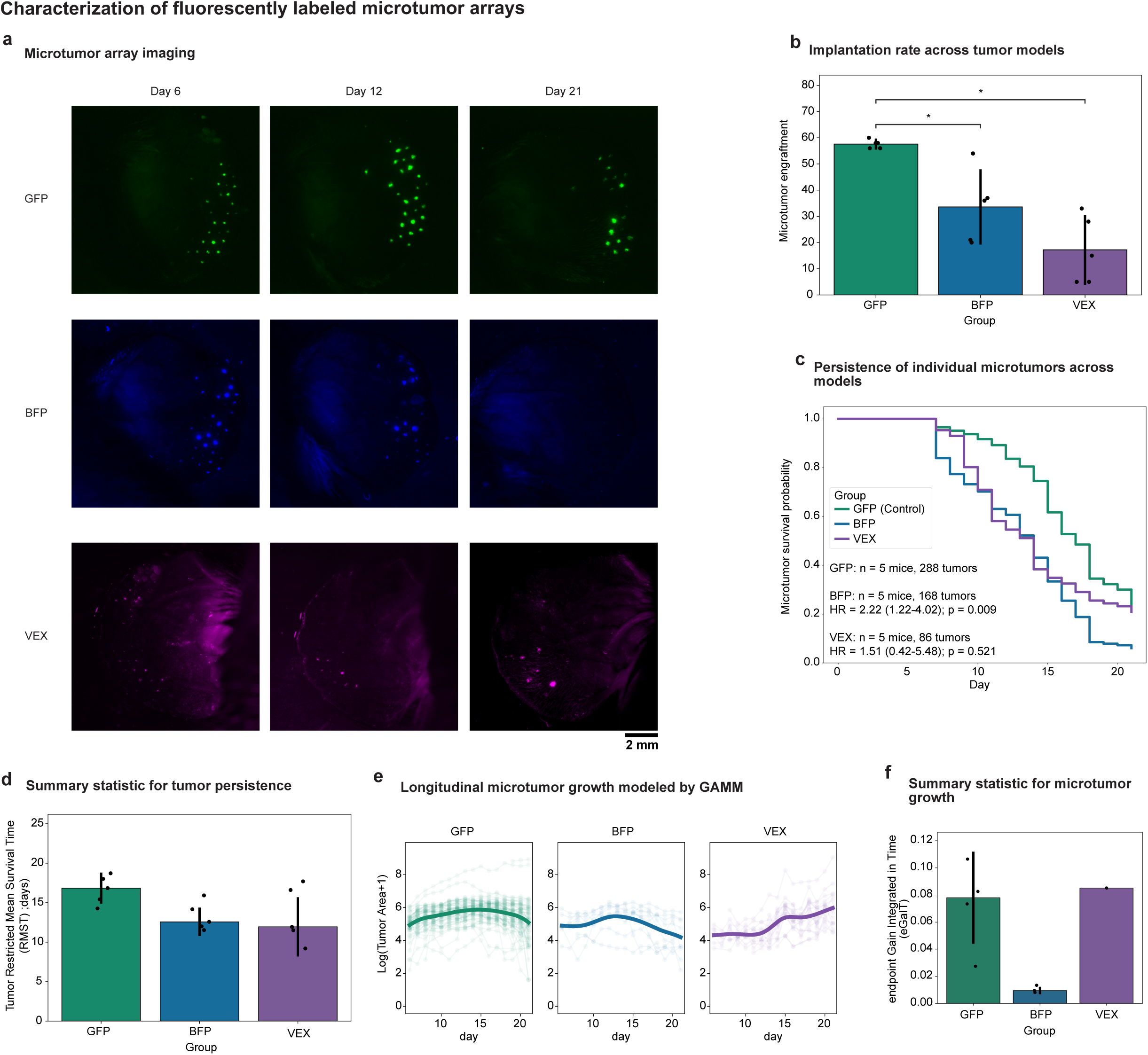
DERM-Array enables longitudinal analysis of tumors expressing eGFP, mTagBFP2, or VEX fluorescent reporters. B16F10 tumor arrays expressing eGFP (GFP), mTagBFP2 (BFP), or VEX fluorescent proteins (n = 5 mice per group) were implanted in WT mice and analyzed for microtumor engraftment, growth, and rejection. **a.** Representative images of tumor arrays on days 6, 12, and 21 post-implantation. Scale bar, 2 mm. **b.** Microtumor engraftment rate per mouse. **c.** Kaplan–Meier curves showing rejection of individual microtumors, with tumors pooled across all mice within each fluorophore cohort. **d.** Restricted mean survival time (RMST) of microtumors per mouse. **e.** Growth trajectories (log-transformed) of persisting microtumors by fluorophore; thin lines represent individual microtumors and bold lines represent the GAMM-fitted summary trajectory for each cohort. **f.** Endpoint Gain Integrated in Time (eGaIT), a cumulative measure of tumor growth, for persisting microtumors per mouse. For panels **b, d**, and **f,** each point represents a single mouse. Bars represent group mean and error bars indicate standard deviation. Statistical analysis was performed using a two-tailed Mann–Whitney U-test; hazard ratios were calculated using Cox Proportional Hazards Model.

**Supplementary Figure 7.**
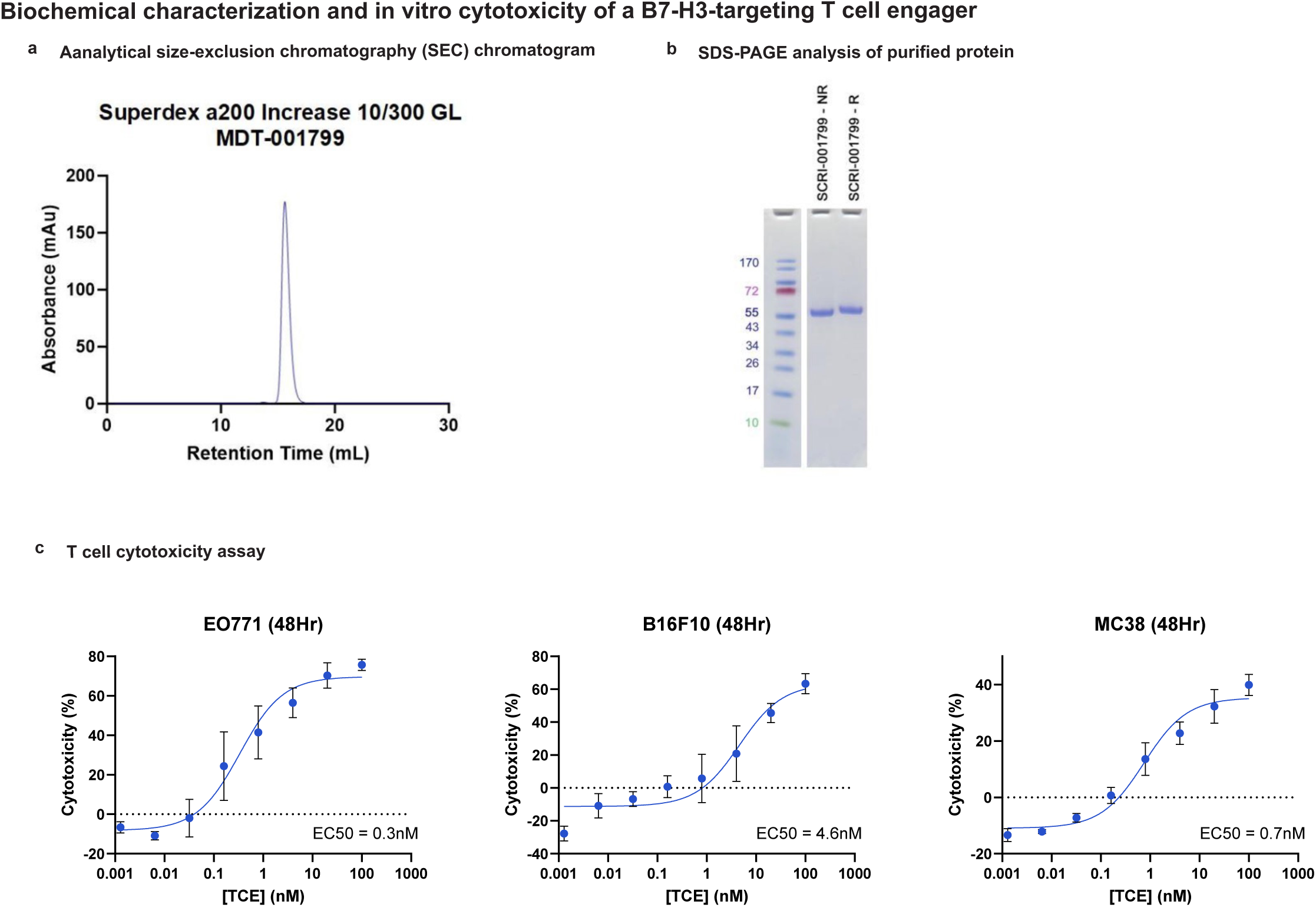
Purified anti-B7-H3/anti-CD3 T cell engager drives dose-dependent tumor cell lysis in vitro. **a.** Representative analytical size-exclusion chromatography (SEC) chromatogram of purified anti-B7-H3/anti-CD3 T cell engager showing a predominant elution peak. SEC calibration standards: 670 kDa (9.3 mL), 158 kDa (12.4 mL), 44 kDa (15.2 mL), 17 kDa (17.5 mL), and 1.3 kDa (20.8 mL). **b.** Coomassie-stained SDS-PAGE analysis of purified anti-B7-H3/anti-CD3 T cell engager under non-reducing (NR) and reducing (R) conditions. **c.** Tumor cells (B16F10, EO771, MC38) expressing eGFP and T cells were co-cultured with varying concentrations of anti-B7-H3/anti-CD3 T cell engager (TCE). Tumor cell lysis at 48 hours relative to baseline. EO771 (left), B16F10 (middle), MC38 (right).

## Supplementary Note 1. Statistical Methods

Here we suggest a statistical analysis framework for interpreting tumor array data.

### Tumor survival

Tracking the survival of individual tumor lesions within an array is an informative readout for many experiments using the DERM-Array model, especially when evaluating therapeutic efficacy. We use Kaplan-Meier survival analysis for visual representation of overall tumor survival dynamics and restricted mean survival time (RMST) for a simple summary statistic at the individual mouse and group levels.

We generate unweighted Kaplan-Meier (KM) curves for all tumors per mouse and per group using KaplanMeierFitter from the lifelines package (**Fig. 2D**) (Davidson-Pilon et al. 2019). For DERM-Array data, KM curves represent tumor survival where each step down in the curve indicates an individual lesion becoming rejected; thus, unlike conventional survival curves, a downward shift reflects increased tumor rejection and a more favorable outcome for the mouse. These curves provide an easily interpretable summary of overall trends in tumor persistence and rejection across individuals and groups.

To summarize survival differences between groups, we used the semi-parametric Cox’s Proportional Hazards model (**Equation 1**), with the baseline hazard (*h*_0_(*t*)) estimated non-parametrically using Breslow’s method using CoxPHFitter from the lifelines package (**Supplementary Fig. 5B, 6C**) (Davidson-Pilon et al. 2019; Royston et al. 2002). We specify individual mouse ID as unique identifiers for clustering covariances; the package applies a robust variance (sandwich) estimator to account for violating assumptions of independence. This model outputs hazard ratios, *p*-values, and 95% confidence intervals that compare experimental groups to a control group. We caution that some experimental designs may result in data with non-proportional hazards (e.g. survival curves cross) and this model may not be ideal in those cases.

For a simple summary statistic that accurately reflects mouse-to-mouse variability, we calculate restricted mean survival time (RMST) for each mouse and then perform a Mann-Whitney U-test to determine significance between groups (**Fig. 2C**). The time threshold, tau (t), is set as the last day of data collection. RMST is a robust measurement for median time to tumor rejection, even when tumor survival never falls below 50% (unlike MST). Further, this complements survival curves by condensing complex survival dynamics into a single mouse-level or group-level summary statistic that represents the area under the KM curve until the endpoint (t).

### Tumor engraftment

Tumor engraftment can vary across mice due to factors such as inconsistencies in laser implantation, baseline differences in host immunity, and cage effects, for example. Assessing tumor engraftment can offer insight into implantation variability and provide context for interpreting results. We quantify total tumor engraftment per mouse and then perform a Mann-Whitney U-test to determine significance between groups (**Fig. 2B**).

### Tumor growth

Tumor growth is a standard measure of treatment response in preclinical mouse tumor studies and can capture changes in persistent tumors (tumors that remain detectable and are not rejected), including slowed growth or partial response, that may be missed by survival analysis alone. We implemented the strategy described in Forrest et al. (2020) which models longitudinal tumor growth using flexible spline-based fits and summarizes each fitted trajectory using endpoint Gain Integrated in Time (eGaIT). In short, individual tumor sizes were log-transformed (**Equation 2**) and plotted over time for persistent tumors at the mouse and/or group level. A generalized additive mixed model (GAMM), implemented with pyGAM (Brummitt et al. 2018), was then fitted to the mean tumor size at each time point (**Fig. 2F**). Next, we calculated eGaIT from the fitted curve to provide an intuitive, biologically interpretable measure of tumor growth, akin to slope in log-linear models (**Fig. 2G**; **Equation 4**; Forrest et al. 2020). Thus, this provides a complementary measure of treatment effect.

## Equations

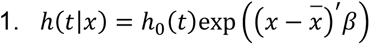

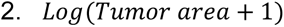

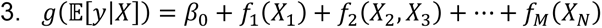

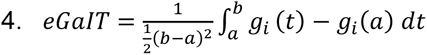

## Supplementary method: DERM-Array protocol

### Animals

#### Mice

Tumor array recipient mice; age 8-10 weeks Immune cell donor mice; age 8-10 weeks

### Materials

#### Depilation

Inhalant anesthesia system with isoflurane (Patterson 07-894-8943) VEET cream (Amazon B001KYPZ4G) Cotton tip applicators (Fisher Scientific 22029683) Kimwipes (Kimtech #34120) Sink water stream

#### Mouse identification

Tattoo ink (Amazon B07HQPPZY6) 27G 1/2in needles (BD 305109)

#### T cell transfer

6 well plate (Corning 3516) 100 µm cell strainer (Fisher Scientific #502274279) Syringe plunger EasySep™ Mouse CD8+ T Cell Isolation Kit (Stem Cell #19853) 15 ml conical tube (VWR 89039-664)

#### Tumor Cell Suspension

Confluent cell culture flasks (∼1.0×10^7^ cells per mouse)

Trypsin or other detachment reagent

FBS-containing media

Phosphate-buffered saline (PBS; 1X)

Ice bucket

#### Anesthesia

1 ml syringes (BD 309659)

27G 1/2in needles (BD 305109)

Ketamine (100 mg/kg; Patterson #07-894-8462)

Xylazine (15 mg/kg; Patterson #07-891-6165). Sterile Saline (as diluent; Patterson #07-892-4348)

Altalube or equivalent ophthalmic ointment (Systane 0065050939)

Heating pad

#### Laser ablation and tumor implantation

1-inch double-sided adhesive tape (Tesa 4970)

Acrylic sheets (1/2×8×8 inches; TAP Plastics) as mounting tray Tweezers, scissors

Charcoal Powder (Millipore Sigma #242276-250G)

Latex-free wedge-style cosmetic sponge (Amazon ASIN B0BNZZQZN2)

XTool F1 Ultra device, controller touchpad, and XCS software (with pre-set design file) Tumor cell suspension (5.0-10×10^6^ cells in 100 µl PBS per mouse ear)

P200 micropipette and tips

#### Longitudinal imaging

Inhalant anesthesia system with induction chamber and nose cone attachment secured to microscope stage

Isoflurane (Patterson 07-894-8943)

1/2-inch double-sided tape (Thomas Scientific #1520CL100P3D)

Epifluorescence stereomicroscope (Leica MF205A) with relevant filters, attachments, software (LASX)

### Day -1: Preparation

#### Depilation (hair removal)

Heavily anesthetize mice with isoflurane.

Apply an even layer of VEET cream to front and back surfaces of both ears with a cotton swab. Return mouse to isoflurane chamber and allow to incubate for 1-3 minutes.

Use a kimwipe to remove most of the product, then rinse mouse ears liberally under a running stream of water to remove all residual product.

Note 1: Residual VEET cream will cause skin irritation and scratching, as well as interfere with tumor growth. Visual inspection and physical texture are reliable indicators – if you see product on the skin or there is a slippery feel to mouse ears, perform an additional rinse.

Note 2: Mice will likely wake up during the rinse step; ensure the mouse is properly restrained with a scruff and work quickly. Avoid getting water in mouse’s airway.

#### Mouse identification

Immobilize mice via isoflurane or firm manual restraint.

Gently extend the paw away from the body and repeatedly insert a needle coated in tattoo ink into the pad surface.

Note: Paw pad tattoos offer up to 17 unique IDs, including no tattoos.

#### T cell isolation and transfer

Harvest spleens from donor mice and place into a 6-well plate with 3 ml cold PBS per well. Keep on ice.

Dissociate spleen by using a syringe plunger back to crush the tissue through a 100 µm cell strainer.

Transfer to a 15 ml conical tube, then spin dissociated splenocytes at 1000 x g for 5 minutes at 4°C. Discard supernatant and resuspend in 1 ml PBS per spleen.

Use the EasySep CD8+ T cell isolation kit according to manufacturer instructions:

Add 20 µl FcR blocker and 50 µl isolation cocktail per spleen sample. Mix and incubate at room temperature for 10 minutes.

Vortex RapidSpheres for 30 seconds, then add 125 µl per spleen sample. Mix and incubate for 5 minutes.

Top up tubes with PBS – up to 5 ml if you have less than 5 spleens, 10 ml if you have 5+ spleens. Mix by pipetting up and down.

Place the tubes onto the EasyEights magnet and incubate for 5 minutes.

Carefully remove the enriched supernatant using a 5 ml serological pipette and transfer to a new tube. Do not disturb the beads or remove tube from magnet.

Count cells. Spin at 1000 x g for 5 minutes, discard supernatant, and resuspend in desired volume according to cell count.

Administer T cells via retro orbital injection. We typically inject 1×10^6^ cells in 100 µl of PBS per mouse.

### Day 0: Implantation

#### Cell Culture

Prior to the day of implantation, grow several large flasks of chosen cell line. A confluent T225 typically contains 5.0-10.0×10^7^ cells; aim for at least 5×10^6^ cells per mouse ear.

Immediately before procedure, remove media, wash with PBS, and detach cells with trypsin (if needed). Neutralize trypsin with FBS-containing media and spin at 500 x g for 5 minutes.

Discard supernatant and resuspend in a small amount of PBS.

Count cells and adjust total volume, re-spinning and resuspending if needed. Each mouse ear will receive 100 µl of cell suspension and most cell lines implant well with 5×10^6^ cells per mouse ear.

#### xTool Set Up

Review XTool safety guidelines (https://support.xtool.com/article/1366). Ensure all institutional regulations are followed regarding laser systems and other safety concerns.

Note: We suggest connecting the device to a vacuum extraction system to reduce smoke emissions.

Power on machine and connect accessories.

Plug in XTool power and turn on machine by pressing the silver button. Confirm red emergency stop button is not depressed.

Ensure the touchpad is plugged into the machine (touchscreen controller port). Connect laptop via USB-C cord (USB port).

Turn on vacuum or other ventilation system.

Open file for microporation arrays in xTool Creative Studio, connect device, and assess calibration.

Confirm accurate calibration by observing the red and blue light dots on the workspace (with the green safety shield up). If dots are precisely overlaid, calibration is acceptable. If not, run the XCS recalibration program or manually adjust using a 3/32 inch/2.5 mm hex screwdriver (manufacturer-provided tool or Allen key).

#### Mouse preparation

Anesthetize mice via intra peritoneal injection of anesthetic mixture (100 mg/kg ketamine and 15mg/kg xylazine per mouse) in sterile saline. Allow mice to become sufficiently anesthetized.

Note: Mice must be kept on heating pad while under anesthesia. Apply ophthalmic ointment to eyes to prevent drying.

Apply a strip of double-sided tape across the middle of a mounting tray (black acrylic 1/2x8x8”). Transfer mice to tray and secure ears to tape by firmly pressing back of each ear flat. 5 mice maximum per tray.

Note: Mounting position determines visibility of ears through laser device software. Try to pull ears upwards and outwards with a moderate amount of tension to maximize visibility while reducing detachment from tape. Mount mice as far apart as possible (at least an inch) to reduce potential for unintended lasing.

Apply charcoal powder to ears using a cosmetic sponge. Dip the sponge into a small pile of charcoal powder, then firmly press onto ears and drag across several times, changing direction as needed, until there is an even layer of powder.

#### Laser ablation

Once mouse and device preparation are complete, place trays in workspace; their bodies should be vertical for best visibility (the tape should be horizontal on your screen).

Position arrays. Refresh the workspace camera view on XCS and align array objects over ears, taking care to cover as much of the visible ear as possible without hitting any fur or eyes.

Note: Do not resize the array at all - this will significantly change the amount of energy applied to the ears.

Once arrays are positioned, click the “Process” button, then check the processing path to confirm that there are no additional objects to be processed. If safe to proceed, process the file and execute via the touchpad.

Note: Ensure green protective cover is fully down, as laser beams are hazardous and can cause damage.

Monitor laser to ensure only ears are lasered and mice continue to be fully sedated. Remove mice from workspace when finished and return to heating pad.

#### Tumor implantation

Gently pipet 100 µL of homogenized tumor cell suspension onto each ear. Use pipet tip to spread meniscus so it covers the whole ear but does not spill onto the tape. Incubate 30-45 minutes.

Gently remove suspension by wicking with a kimwipe. Remove mice from tray and fold ears to protect suspension. Monitor animals continuously until they wake up.

### Days 1-21: Additional steps and data collection

#### Post-implantation care

For the first three days post-implantation, generously apply petroleum jelly to mouse ears with a cotton swab. Apply as needed until desiccation and scabbing resolve.

#### Longitudinal imaging

Anesthetize mice using an isoflurane system and apply ophthalmic ointment to eyes. Continuously monitor mouse status throughout imaging.

Load microscope settings and set zoom based on experimental needs; 10X zoom is sufficient for tumor growth analysis, while 30X zoom is necessary for immune cell analysis.

Cut a piece of double-sided tape about 3-4 inches to have enough room to stick the ears down. Place just below the nose cone on the microscope stage.

When mice are sufficiently anesthetized, remove a mouse from induction chamber, transfer to microscope stage, and place mouse face-up in nose cone. Carefully extend ears and secure flat to tape.

Locate ears on live view and set a region of interest, if needed. Image both ears with a 10X objective if capturing tumors only or a 30X objective if imaging tumors and T cells.

To remove mice from stage, peel one ear up and carefully roll the mouse off the tape in the direction of the other ear. Place mouse onto a paper towel in the home cage for recovery. Replace tape and repeat with remaining mice.

